# Genomic analyses of population structure reveal metabolism as a primary driver of local adaptation in *Daphnia pulex*

**DOI:** 10.1101/807123

**Authors:** Takahiro Maruki, Zhiqiang Ye, Michael Lynch

**Affiliations:** Center for Mechanisms of Evolution, Biodesign Institute, Arizona State University, Tempe, AZ 85287-7701

**Keywords:** population structure, local adaptation, population genomics, *Daphnia pulex*

## Abstract

Elucidating population structure is important for understanding evolutionary features of an organism. In the freshwater microcrustacean *Daphnia pulex*, an emerging model system in evolutionary genomics, previous studies using a small number of molecular markers indicated that genetic differentiation among populations is high. However, the dispersal ability of *D. pulex* is potentially high, and evolutionary forces shaping genetic differentiation among populations are not understood well. In this study, we carried out genomic analyses using high-throughput sequencing to investigate the population structure of *D. pulex*. We analyzed 10 temporary-pond populations widely distributed across the midwestern United States, with each sample consisting of 71 to 93 sexually reproducing individuals. The populations are generally in Hardy-Weinberg equilibrium and have relatively large effective sizes. The genetic differentiation among the populations is moderate and positively correlated with geographic distance. To find outlier regions showing significantly high or low genetic differentiation, we carried out a sliding-window analysis of the differentiation estimates using the bootstrap. Genes with significantly high genetic differentiation show striking enrichment of gene ontology terms involved in food digestion, suggesting that differences in food quality and/or quantity among populations play a primary role in driving local adaptation of *D. pulex*.

---

Species generally consist of metapopulations of demes connected by some amounts of gene flow. As individual populations may experience different ecologies, identifying population similarities and differences shaped by geographic or ecological factors is important for understanding mechanisms of evolution. In the freshwater microcrustacean *Daphnia pulex*, pond populations have well-defined boundaries, providing excellent opportunities for studying their structure. Previous studies on the population structure in *Daphnia pulex* used allozymes, microsatellites, or mitochondrial DNA as molecular markers (Crease *et al*. 1990; Lynch and Crease 1990; Innes 1991; Lynch and Spitze 1994; Morgan *et al*. 2001; Allen *et al*. 2010). Most of these studies (Crease *et al*. 1990; Lynch and Crease 1990; Lynch and Spitze 1994; Morgan *et al*. 2001; Allen *et al*. 2010) suggested that the genetic differentiation among populations is high. On the other hand, one study (Innes 1991) using six allozyme loci indicated moderate genetic differentiation among populations.

When stressed, *Daphnia* can produce resting eggs surrounded by protective membranes (Hebert 1978). These ephippia are resistant to desiccation and digestion, have well-developed spines along the dorsal margin, which facilitate their attachment to other animals and therefore enable passive dispersal. Thus, although the dispersal ability of live *Daphnia* is limited, their resting eggs can be transported to other areas by wind, flowing water, or animals such as birds (Havel and Shurin 2004; Figuerola *et al*. 2005). Furthermore, resting eggs deposited in pond/lake sediments can remain dormant for long periods (Brendonck and De Meester 2003), enabling *Daphnia* dispersal over time. Given the potentially high dispersal ability of *Daphnia* enabled by these mechanisms, the high genetic differentiation found in previous population-genetic studies of *D. pulex* is puzzling.

Due to the lack of genomic data, population-genetic studies in *D. pulex* have been limited until recently, and were based on a small number of molecular markers. When a small number of molecular markers are used, the estimate of genetic differentiation among populations can vary from study to study, and it is difficult to infer the evolutionary forces shaping the genetic differentiation. The recent publications of genomic reference sequences of *D. pulex* (Colbourne *et al*. 2011; Ye *et al*. 2017) and high-throughput sequencing technologies have changed the situation, enabling whole-genome population-genetic analyses of many individuals based on single nucleotide polymorphisms (SNPs) (Lynch *et al*. 2017).

In this study, we carry out population-genomic analyses of 10 *D. pulex* populations to investigate the evolutionary forces shaping the genomic differentiation among the populations. The populations are from temporary ponds, which dry up annually, and are widely distributed across the midwestern United States. We sequenced 96 clones from each of the populations. By sequencing randomly across the entire genome, our study enables unbiased estimation of genetic differentiation among the populations based on genomic SNPs.

Our poulation-genomic analyses have significant advantages over the previous population-genetic analyses in *D. pulex*. SNPs provide the finest molecular analyses, and by analyzing huge numbers of SNPs across the genome, we can find signatures of natural selection in much less biased ways than previous studies. As population-genomic studies identify putatively selected regions without prior knowledge, they may identify functionally important genetic regions difficult to find using other approaches. Finding targets of natural selection is important for studying population structure (Luikart *et al*. 2003). Most population-genetic models used for estimating parameters of population demography/structure assume neutral evolution. By identifying putative targets of selection and removing them beforehand, we can minimize the confounding effect of natural selection on the inference of population demography/structure.

Our results indicate that genetic differentiation among *D. pulex* populations is moderate and positively correlated with geographic distance. In addition, we identify intriguing signatures of local adaptation likely shaped by nutritional differences among populations. Together with comparisons of divergence/heterozygosity estimates at replacement and silent sites in protein-coding sequences, our genomic analyses suggest the importance of nutrition in adaptations of *Daphnia* species.

## MATERIALS AND METHODS

### Sample preparation and sequencing

We randomly collected *Daphnia pulex* individuals from 10 temporary ponds in the spring of 2013 or 2014 across the midwestern United States (TABLE 1). As in Lynch *et al*. (2017), to maximize the likelihood that each individual would originate from a unique resting egg, we collected adult individuals soon after hatchlings were found and before substantial reproduction had occurred. We clonally expanded the individual isolates, extracted DNA, and sequenced DNA of 96 isolates per population, tagging each isolate by unique oligomer barcodes, with paired-end short reads using Illumina technologies as previously described (Lynch *et al*. 2017).

**TABLE 1.** Information on the analyzed populations. Sample size denotes the number of analyzed clones. Population coverage denotes the chosen range (in square brackets) and mean of the population-wide sum of the depth of coverage of the analyzed nucleotide reads over the clones.

| Population | (Latitude, Longitude) | Sample Size | Population Coverage |
| --- | --- | --- | --- |
| Busey Pond (BUS) | (40.13, -88.21) | 89 | [70, 1050], 350.9 |
| Chequamegon (CHQ) | (46.66, -90.85) | 93 | [500, 2200], 1323.2 |
| Eloise Butler Pond (EB) | (44.98, -93.32) | 82 | [225, 1050], 578.2 |
| Kickapond (KAP) | (40.12, -87.74) | 81 | [800, 2400], 1405.2 |
| Long Point Pond A (LPA) | (42.69, -80.42) | 88 | [400, 2250], 1105.6 |
| Long Point Pond B (LPB) | (42.68, -80.45) | 84 | [400, 1900], 1026.0 |
| North Flatley (NFL) | (39.90 -84.92) | 89 | [700, 2200], 1442.4 |
| Portland Arch (PA) | (40.21, -87.33) | 75 | [150, 1900], 670.8 |
| Pond of the Village Idiot (POV) | (42.75, -85.35) | 71 | [500, 2250], 1093.2 |
| Textile Road (TEX) | (42.20, -83.60) | 72 | [250, 1800], 631.5 |
| **Total** | **NA** | **824** | **[3995, 19000], 9282.2** |

### Data preparation

From the FASTQ files of sequence reads, we prepared nucleotide-read quartets (counts of A, C, G, and T) necessary for the population-genomic analyses, taking various filtering steps to process high-quality data. First, we trimmed adapter sequences from the sequence reads by applying Trimmomatic (version 0.36) (Bolger *et al*. 2014) to the FASTQ files. Then, we mapped the adapter-trimmed sequence reads to PA42 (version 3.0) (Ye *et al*. 2017), using Novoalign (version 3.02.11) (http://www.novocraft.com/) with the “-r None” option to exclude reads mapping to more than one location. To reduce the computational time taken for mapping, we split the FASTQ files using NGSUtils (version 0.5.9) (Breese and Liu 2013), mapped them separately to PA42, and combined the resulting SAM files using Picard (version 2.8.1) (http://broadinstitute.github.io/picard/). We converted the SAM files of the mapped sequence reads to BAM files using Samtools (version 1.3.1) (Li *et al*. 2009). We marked duplicates and locally realigned indels in the sequence reads stored in the BAM files using GATK (version 3.4-0) (McKenna *et al*. 2010; DePristo *et al*. 2011; Van der Auwera *et al*. 2013) and Picard. In addition, we clipped overlapping read pairs by applying BamUtil (version 1.0.13) (http://genome.sph.umich.edu/wiki/BamUtil) to the BAM files. We made mpileup files from the processed BAM files using Samtools. We then made a file of nucleotide-read quartets of all individuals, called a pro file, in each of the populations using the proview command of MAPGD (version 0.4.26) (https://github.com/LynchLab/MAPGD).

We estimated the allele and genotype frequencies in each population by the maximum-likelihood (ML) method of Maruki and Lynch (2015), using the allele command of MAPGD. To minimize analyzing sites with mismapped reads, we eliminated sites that showed poor fit to the estimated parameters from subsequent analyses by the goodness-of-fit test (Ackerman *et al*. 2017), setting the minimum number of individuals with poor fit to the data required for eliminating the site at four. Poor fit of the estimated parameters to the observed data at a site can happen, for example, when some sequence reads mapping to the site originate from paralogs not found in the genome assembly.

To further refine the data, we removed clones with the mean coverage over sites less than three from subsequent analyses. We also removed clones with the sum of the goodness-of-fit values across the genome less than -0.4, which can happen, for example, when a clone is contaminated with another clone or belongs to another species. In addition, to avoid including closely related individuals, we estimated the pairwise relatedness of the clones using the relatedness command of MAPGD (Ackerman *et al*. 2017) and kept only the clone with the highest coverage in each of the clusters of clones with the relatedness estimates *≥* 0.125. Furthermore, to avoid including obligately asexual clones in the analyses, we examined the fractions of sites containing asexual-specific alleles at the asexual markers (Tucker *et al*. 2013; Xu *et al*. 2015) in each of the clones, and excluded those with the fraction *≥* 0.03. The resulting total number of analyzed clones in each population is shown in TABLE 1. In the subsequent analyses, we removed sites involved in putatively repetitive regions identified by RepeatMasker (version 4.0.5) (http://www.repeatmasker.org/) with the RepeatMasker library (Jurka *et al*. 2005) made on August 7, 2015. Furthermore, we examined the genomic distribution of the population coverage (sum of the depths of coverage over the clones), and set the minimum and maximum population-coverage cut-offs to avoid analyzing problematic sites (TABLE 1). Specifically, we chose the local minimum closest to the mode of the representative distribution and a value at the edge of the representative distribution to set the minimum and maximum population-coverage cut-offs, respectively. In addition, we removed sites with ML error-rate estimates *>* 0.01.

In summary, our analyses are conservative with respect to both individuals and sites. We removed individuals that are potentially contaminated in the laboratory, have close relatives in the population sample, or are obligatory asexual. We removed sites that are potentially involved in problems with paralogs or other mapping issues.

### Estimation of the number of alleles

To estimate the number of alleles at each site, we applied the high-coverage genotype caller (HGC) (Maruki and Lynch 2017) to the data in each population before filtering sites by the goodness-of-fit test. We set the minimum coverage required for calling genotypes at six to avoid the false-positive detection of alleles (Maruki and Lynch 2017). Because most downstream analyses assume biallelic polymorphisms, we excluded multi-allelic sites in each population from subsequent analyses, setting the significance of called genotypes at the 5% level. We note that polymorphic sites can still contain three or four alleles at the metapopulation level and we analyzed such sites.

### Estimation of Wright’s fixation indices

We estimated Wright’s fixation indices (1951) from the genotype-frequency estimates at SNP sites in each population, which were predetermined by the genotype-frequency estimator (Maruki and Lynch 2015). To avoid analyzing false alleles resulting from sequencing errors, we restricted these analyses to sites significantly polymorphic in at least one of the populations at the 5% level. We estimated the fixation indices in the framework of Weir and Cockerham (1984), where *F_IT_* (inbreeding coefficient in the total population), *F_ST_* (measure of genetic differentiation), and *F_IS_* (mean inbreeding coefficient within populations) are determined from the genotype frequency estimates (Weir 1996).

The method of Weir and Cockerham has been designed for estimating fixation indices from accurate genotypes with no missing data. However, in data generated by high-throughput sequencing, depths of coverage vary among sites, individuals, and chromosomes within diploid individuals, due to random sequencing. Therefore, adjustments are needed to take the variability of the sample size into account. We did so by estimating the effective number of sampled individuals (Maruki and Lynch 2015) at each site in each deme. The effective number of sampled individuals *n_ei_* at a site in deme *i* is the expected number of individuals for which both chromosomes are sequenced at least once, and is estimated as

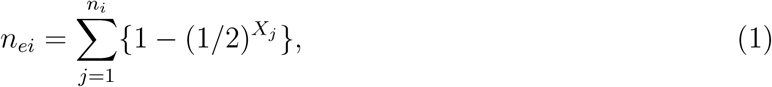

where *n_i_* and *X_j_* are the number of sampled individuals in deme *i* and depth of coverage in individual *j*, respectively. We required that *n_ei_* be at least 10 in each deme, estimating the fixation indices using data from demes with *n_ei_ ≥* 10. Letting *r_e_* denote the number of demes with *n_ei_ ≥* 10, we estimated the fixation indices by substituting *n_ei_* and *r_e_* for the number of sampled individuals in deme *i* and number of demes, respectively, in the equations by Weir and Cockerham.

### Analyses of pairwise *F_ST_* estimates

To examine the relationship between geography and genetic differentiation among the populations, we built a neighbor-joining tree (Saitou and Nei 1987) based on the mean pairwise *F_ST_* estimates, using the R package ape (version 5.0) (Paradis *et al*. 2004). We also made a scatter plot of the geographic distance and mean pairwise *F_ST_* estimates between populations at silent and replacement sites in protein coding sequences, inferring the former from the coordinates of the populations using the R package geosphere (version 1.5-7) (Hijmans 2017).

### Sliding-window analyses of the fixation index estimates

Because the fixation index estimates measured at individual SNP sites are highly variable, we carried out sliding-window analyses of the estimates to examine their spatial patterns along the scaffolds. Each of the windows contained a fixed number of 101 SNPs. To enable finer description of the spatial patterns, we calculated the weighted mean of the estimates in each of the windows with the weight given by 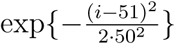, where *i* is a SNP index (*i* = 1, 2, *· · ·,* 101), so that SNPs closer to the middle SNP contribute more to the mean (Hohenlohe *et al*. 2010). 21 SNPs were overlapping between adjacent windows in our analyses.

### Statistical identification of fixation-index outliers

To statistically identify fixation-index outliers, we carried out bootstrap analyses (Efron and Tibshirani 1993). Specifically, we determined the top and bottom critical values for the mean fixation-index estimates at the 5% significance level from their genome-wide distribution based on bootstrap replications. In each bootstrap replication, we randomly sampled 101 fixation-index estimates from genomic SNPs and calculated their weighted means in the way described above. We carried out a total of 10^9^ such bootstrap replications. To take the multiple-testing problem into account, we applied the Bonferroni method by dividing the significance level by the total number of windows in the genome.

### Estimation of *X^T^ X*

Because we analyze multiple populations with different degrees of genetic differentiation among different population pairs, we also estimated *X^T^ X* (Günther and Coop 2013) at the SNP sites. This *F_ST_* analog quantifies the genetic differentiation among populations taking its heterogeneity among different population pairs into account. We used Bayenv2.0 (Günther and Coop 2013) to estimate *X^T^ X*. We prepared the input file of allele counts using the method of Maruki and Lynch (2015). Because the software requires the allele-count data in all of the analyzed populations with two segregating alleles in the total population, we analyzed only sites with such data. We estimated the covariance matrix of the allele-frequency estimates based on 100,000 Markov chain Monte Carlo (MCMC) iterations. We ran 10,000 MCMC iterations to estimate *X^T^ X* at each site, which we found to yield estimates very similar to those based on 100,000 iterations on the largest scaffold (results not shown). We carried out sliding-window and bootstrap analyses to examine the spatial pattern of the *X^T^ X* estimates and identify the outliers, respectively, in ways similar to those for the fixation-index estimates.

### Enrichment analysis of gene ontology terms in the outlier genes

To infer the functions of the *F_ST_* outlier genes, we examined their gene ontology (GO) terms (Ashburner *et al*. 2000) using Blast2GO (Conesa *et al*. 2005). Specifically, we applied BLAST (Altschul *et al*. 1990; Camacho *et al*. 2009) and InterProScan (Jones *et al*. 2014) to the amino-acid sequences of the genes in the PA42 reference genome (Ye *et al*. 2017), mapped them to the GO terms, and annotated them applying the ANNEX augmentation (Myhre *et al*. 2006). To find GO terms enriched in each type of the outlier genes, we carried out the enrichment analysis of the GO terms using Fisher’s exact test accounting for the multiple-testing problem (Al-Shahrour *et al*. 2004) and applying Go-Slim to the GO terms to reduce the redundancy among the terms.

### Comparison of the differentiation estimates at replacement and silent sites

To infer functional differences underlying natural selection shaping the genetic differentiation among the populations, we compared the mean differentiation estimates over zero-fold and four-fold redundant sites in each gene. We used the two-tailed Z-test to identify genes showing an excess or a deficit of amino-acid altering genetic differentiation. To account for the multiple-testing problem, we calculated *q* values (Storey and Tibshirani 2003), which are expected fractions of false-positive tests among significant tests, from the *p* values in the Z-tests using the R package qvalue.

### *d_N_/d_S_* analysis

To identify putative targets of natural selection involved in functional differences of proteins, we carried out a *d_N_/d_S_* analysis in each of the protein-coding sequences, where *d_N_* and *d_S_* are the mean between-species divergence over populations per replacement and silent sites, respectively. We used a draft genome assembly of *D. obtusa* based on hybrid data generated by Illumina short-read and PacBio long-read sequencing to estimate the between-species genetic divergence. Specifically, we aligned the *obtusa* genome sequence to the *pulex* genome sequence (PA42 version 3.0) using LAST (version 912) (Kielbasa *et al*. 2011). The *obtusa* genome sequence is useful for the *d_N_/d_S_* analysis because of its moderate silent-site divergence from the *pulex* genome sequence (the mean = 0.1090 with the standard error = 0.0005) (Lynch *et al*. 2017). We estimated the divergence only at sites where the nucleotide identity (A, C, G, or T) of both species is known without being involved in repetitive regions. In addition, we required the sites to meet the population-coverage cut-off values and have the error rate estimate *≤* 0.01 in each population. We estimated the divergence as the frequency of the *pulex* nucleotides in the population differing from the *obtusa* nucleotide in each population. To minimize the ambiguity of the analysis, we calculated the divergence estimates at replacement and silent sites as those at zero-fold and four-fold redundant sites, respectively. We calculated the mean of the divergence estimates over zero-fold and four-fold redundant sites, respectively, in each gene in each population. Then, for each gene, we calculated the mean and variance of the mean divergence estimates per site over populations. We estimated *d_N_/d_S_* as a ratio of the mean divergence estimates over populations per zero-fold and four-fold redundant sites.

### *π_N_/π_S_* analysis

To infer types of natural selection involved in functional differences of proteins, we also carried out the *π_N_/π_S_* analysis, where *π_N_* and *π_S_* are the mean heterozygosity over populations per replacement and silent sites, respectively, in a way similar to that in the *d_N_/d_S_* analysis. We also carried out an analogous analysis among the populations. Specifically, we calculated the heterozygosity estimates among populations in the framework of Weir and Cockerham (1984) at zero-fold and four-fold redundant sites. We call the mean heterozygosity estimates among populations over zero-fold and four-fold redundant sites *φ_N_* and *φ_S_*, respectively.

### Identification of candidate genes under natural selection

To identify candidate genes under natural selection, we calculated analogs of the neutrality index (NI) (Rand and Kann 1996) and direction of selection (DoS) (Stoletzhki and Eyre-Walker 2011) using the mean divergence/heterozygosity estimates per site over populations in each gene. We calculated the variance of the DoS and NI estimates using the delta method (APPENDIX). To examine the significance of the *d_N_* estimate with respect to the *d_S_* estimate, we calculated a *z* score for the *d_N_ − d_S_* estimate from the mean and standard error of the *d_N_* and *d_S_* estimates. We calculated a *z* score from the mean and standard error of the NI or DoS estimate. To identify candidate genes under natural selection, accounting for simultaneous multiple tests, we calculated *q* values (Storey and Tibshirani 2003) from *p* values in two-tailed *z*-tests using the R package qvalue. We identified candidate genes under natural selection as those with *q <* 0.05 in at least two of the *d_N_ −d_S_*, NI, and DoS estimates.

### Inference of natural selection shaping the functional changes in the genome

To infer dominant types of natural selection shaping the functional differences in the protein-coding sequences in the genome, we calculated the means of the *d_N_/d_S_*, *φ_N_/φ_S_*, *π_N_/π_S_*, neutrality index (NI), and direction of selection (DoS) estimates over genes. Here, we factored out the confounding effects of polymorphisms within and among populations on divergence estimates by subtracting heterozygosity estimates per site within and among populations from divergence estimates per site in each gene separately, and call the adjusted divergence estimates at replacement and silent sites *d′_N_* and *d′_S_*, respectively. That is,

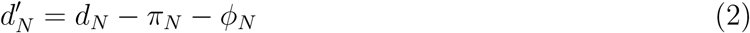

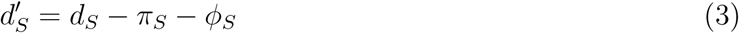

We calculated the means of the *d′_N_/d′_S_*, NI, and DoS estimates only over genes with non-negative *d′_N_* and *d′_S_* estimates. Because these ratios take extreme values when the denominator is close to zero, we excluded genes with estimates greater than five from the calculations of the means over genes.

### Data availability

The FASTQ files of the raw sequencing data are publicly available *via* NCBI Sequence Read Archive (accession number SRP155055). The FASTA file of the draft genome assembly of *D. obtusa* is available upon request. Figure S1 shows the relationship between the mean heterozygosity within populations *π_S_* and *F_ST_* estimates in protein-coding sequences. Figure S2 shows the relationship between the heterozygosity in the total population *π_T_* and *F_ST_* estimates in protein-coding sequences. Table S1 is a list of top *F_IS_* outliers. Table S2 is a list of bottom *F_IS_* outliers. Table S3 is a list of top *F_ST_* outliers. Table S4 is a list of bottom *F_ST_* outliers. Table S5 shows gene ontology terms enriched in the *F_ST_* outlier genes. Table S6 shows gene ontology terms before applying Go-Slim enriched in the top *F_ST_* outlier genes. Table S7 is a list of top *X^T^ X* outliers. Table S8 is a list of bottom *X^T^X* outliers. Table S9 shows gene ontology terms enriched in the *X^T^ X* outlier genes. Table S10 is a list of genes showing a significant excess of amino-acid altering differentiation estimates among populations (*q <* 0.05). Table S11 is a list of genes showing a significant deficit of amino-acid altering differentiation estimates among populations (*q <* 0.05). Table S12 is a list of candidate genes under positive selection (*q <* 0.05). Table S13 shows gene ontology terms enriched in the candidate genes under positive selection. Table S14 is a list of candidate genes under purifying selection (*q <* 0.05). Table S15 shows gene ontology terms enriched in the candidate genes under purifying selection.

## RESULTS

To investigate patterns of polymorphisms in the *Daphnia pulex* populations, we first examined within-population genetic variation. The majority (*>* 95%) of the analyzed sites are monomorphic for the same allele in all of the populations (Figure 1). Among polymorphic sites, the majority (*>* 97%) contain just two nucleotides per site. Tri- and tetra-alleleic sites are generally rare. The mean heterozygosity estimate across genomic sites is similar among the populations (the mean = 0.0078 with the standard error = 0.0004), although the CHQ population has a somewhat lower estimate, suggesting a lower effective population size than in other populations (TABLE 2).

**Figure 1.**
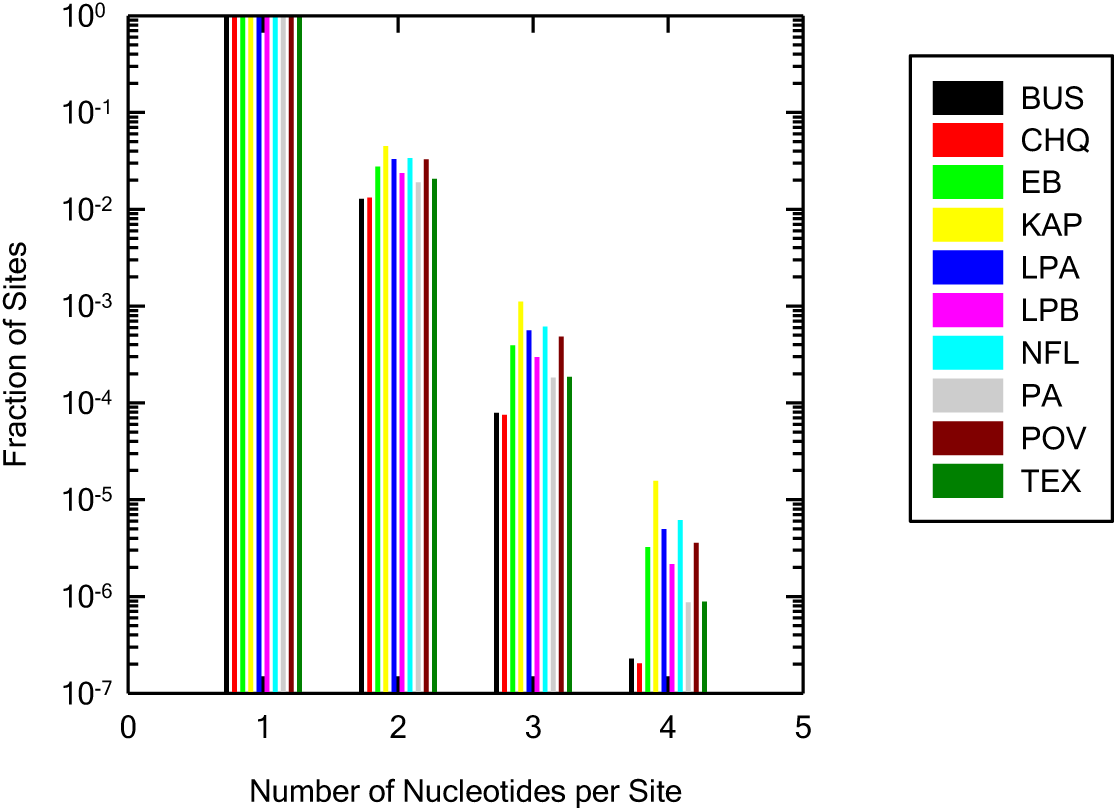
Distribution of the number of nucleotides per site in each population. Statistical significance of the called genotypes is set at the 5% level. In general, tri- and tetra-allelic sites are rare.

**TABLE 2.**
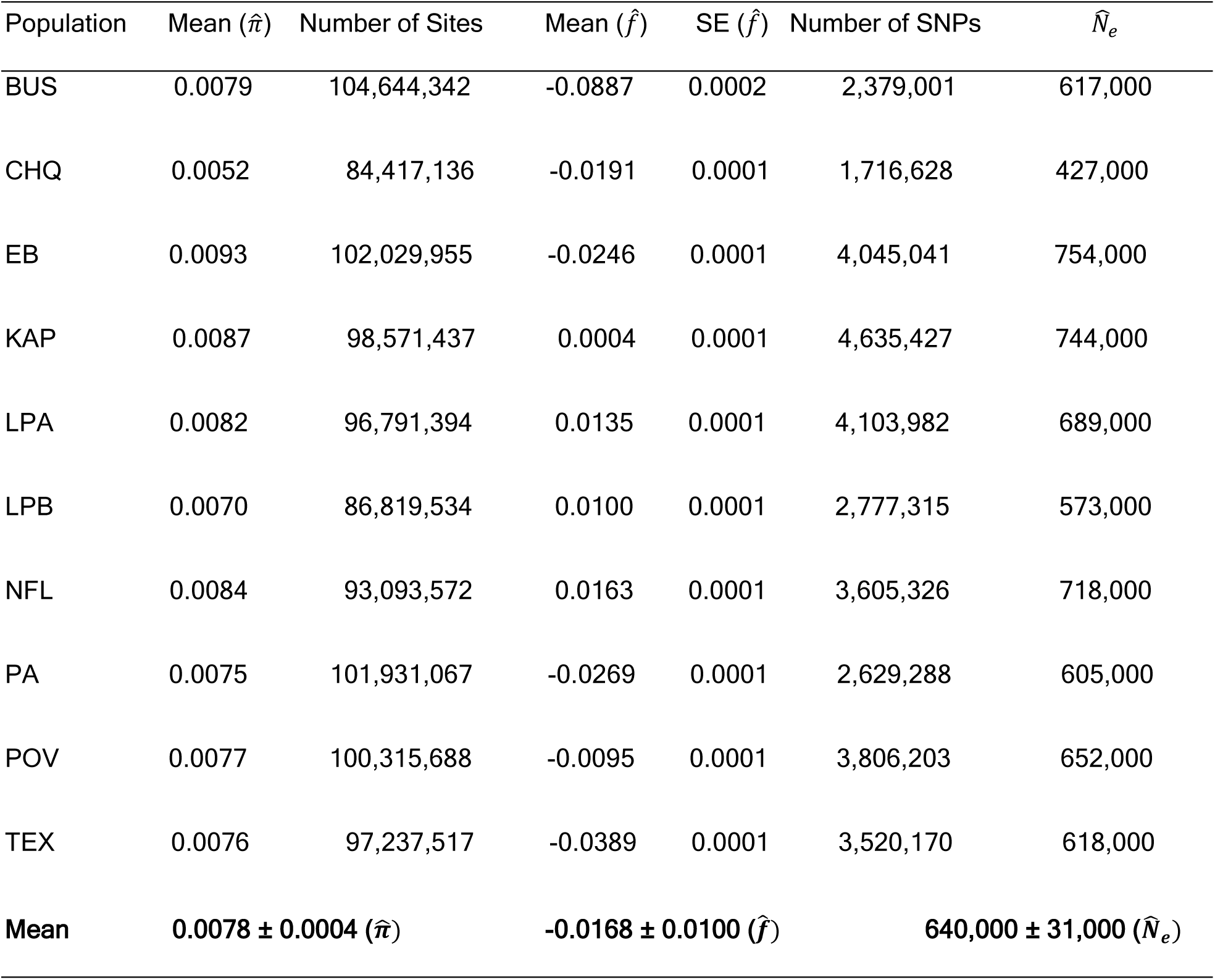
Summary of within-population genetic variation. Mean and standard error (SE) of the heterozygosity and inbreeding coefficient estimates, 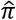 and 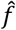, are shown. 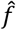 is conditioned on significant polymorphisms at the 5% level. Tri- and tetra-allelic sites are excluded from the calculations. The standard error of the mean heterozygosity estimate 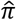 is 0.000005 in all populations. Effective population size estimate 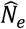, which was estimated using the mutation rate estimate from Keith *et al*. (2016) and mean heterozygosity estimates at silent and restricted intron (internal positions 8 to 34 from both ends; Lynch *et al*. 2017) sites, is also shown. The grand mean and SE of 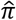, 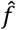, and 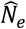 are also shown.

To infer the average effect of natural selection, we examined the mean heterozygosity estimate at sites in different functional categories (Figure 2). The heterozygosity estimate is the highest at silent sites followed by restricted intron (internal positions 8 to 34 from both ends; Lynch *et al*. 2017) sites. It is the lowest at amino-acid replacement sites. Intergenic and UTR sites have intermediate estimates compared to these extremes. These observations are consistent with our previous finding that heterozygosity estimates are lower at functionally more important sites (Lynch *et al*. 2017), indicating that purifying selection decreases heterozygosity estimates.

**Figure 2.**
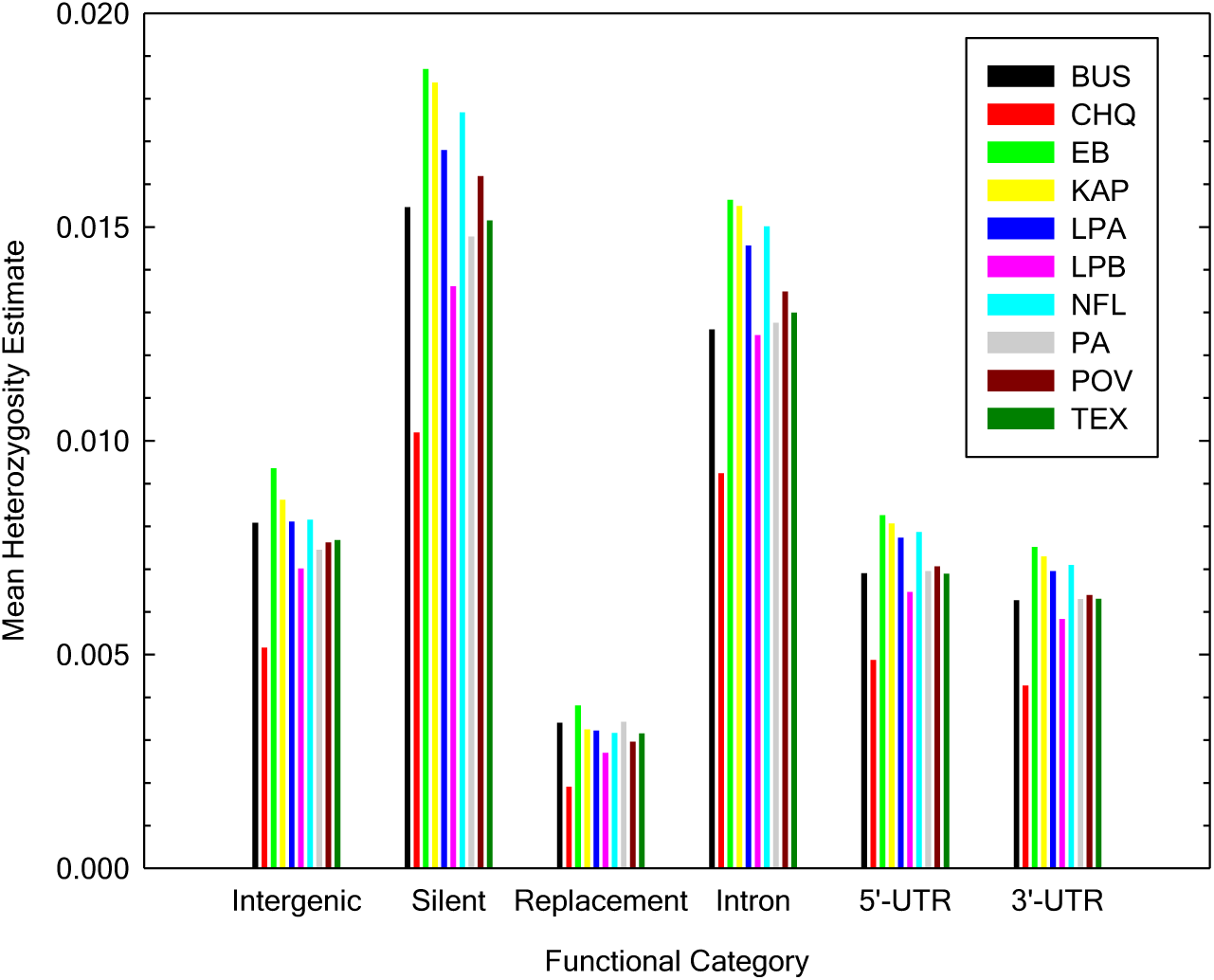
Heterozygosity estimates at sites in different functional categories. Intergenic sites are outside of untranslated regions (UTR), exons, and introns. Silent, replacement, and intron sites are, respectively four-fold redundant, zero-fold redundant and restricted intron (internal positions 8 to 34 from both ends; Lynch *et al*. 2017) sites. Tri- and tetra-allelic sites are excluded from the calculation. The standard errors of the mean are generally small (6.2×10^-6^ to 4.7×10^-5^).

Although the mean inbreeding-coefficient estimate is somewhat negative in the BUS population, it is generally close to zero in the majority of the populations, indicating that they are in Hardy-Weinberg equilibrium in accordance with earlier allozyme estimates (Lynch 1983; Lynch and Spitze 1994) (TABLE 2). The negative mean inbreeding-coefficient estimate in BUS is consistent with Allen *et al*.’s (2010) finding of a negative value (-0.18) in this population using microsatellites. Using the mutation rate estimate *u* = 5.69 *×* 10*^−^*^9^ per site per generation from Keith *et al*. (2016) and the means of the heterozygosity estimates *π* at silent and restricted intron sites, which we found to be essentially under neutral evolution in the analysis of one of the populations (Lynch *et al*. 2017), we estimated the effective size *N_e_* of each population by equating *π* with 4*N_e_u* (TABLE 2). The estimated effective population sizes of the analyzed populations are relatively large and similar to each other (the mean = 640,000 with the standard error = 31,000). Although such estimates assume stable *N_e_*, as described in Lynch *et al*. (in preparation), historical *N_e_* estimates in the study populations are relatively stable and their harmonic means agree reasonably well with the *N_e_* estimates here.

To investigate the genetic structure of the study populations, we estimated Wright’s (1951) fixation indices. The mean of the *F_ST_* estimates is 0.13, indicating moderate differentiation among the populations (TABLE 3). The mean of the *F_IS_* estimates is essentially zero, supporting that within-population polymorphisms are near Hardy-Weinberg equilibrium. Consistent with this, the mean of the *F_IT_* estimates is very close to that of the *F_ST_* estimates. The modes of the genomic *F_IS_* and *F_ST_* estimates are found around zero (Figures 3A and 3B). The median of the *F_IS_* estimates (-0.01) is close to zero, and the median of *F_ST_* estimates (0.08) indicates moderate genetic differentiation among study populations. To infer the influence of natural selection on *F_ST_* estimates, we examined the *F_ST_* estimates at sites in different functional categories (TABLE 4). *F_ST_* estimates are the highest at restricted intron and silent sites and lowest at replacement sites. They are intermediate at intergenic and UTR sites compared to the extremes. The observation here is consistent with previous studies reporting lower *F_ST_* estimates at sites under stronger functional constraints (Barreiro *et al*. 2008; Maruki *et al*. 2012).

**Figure 3.**
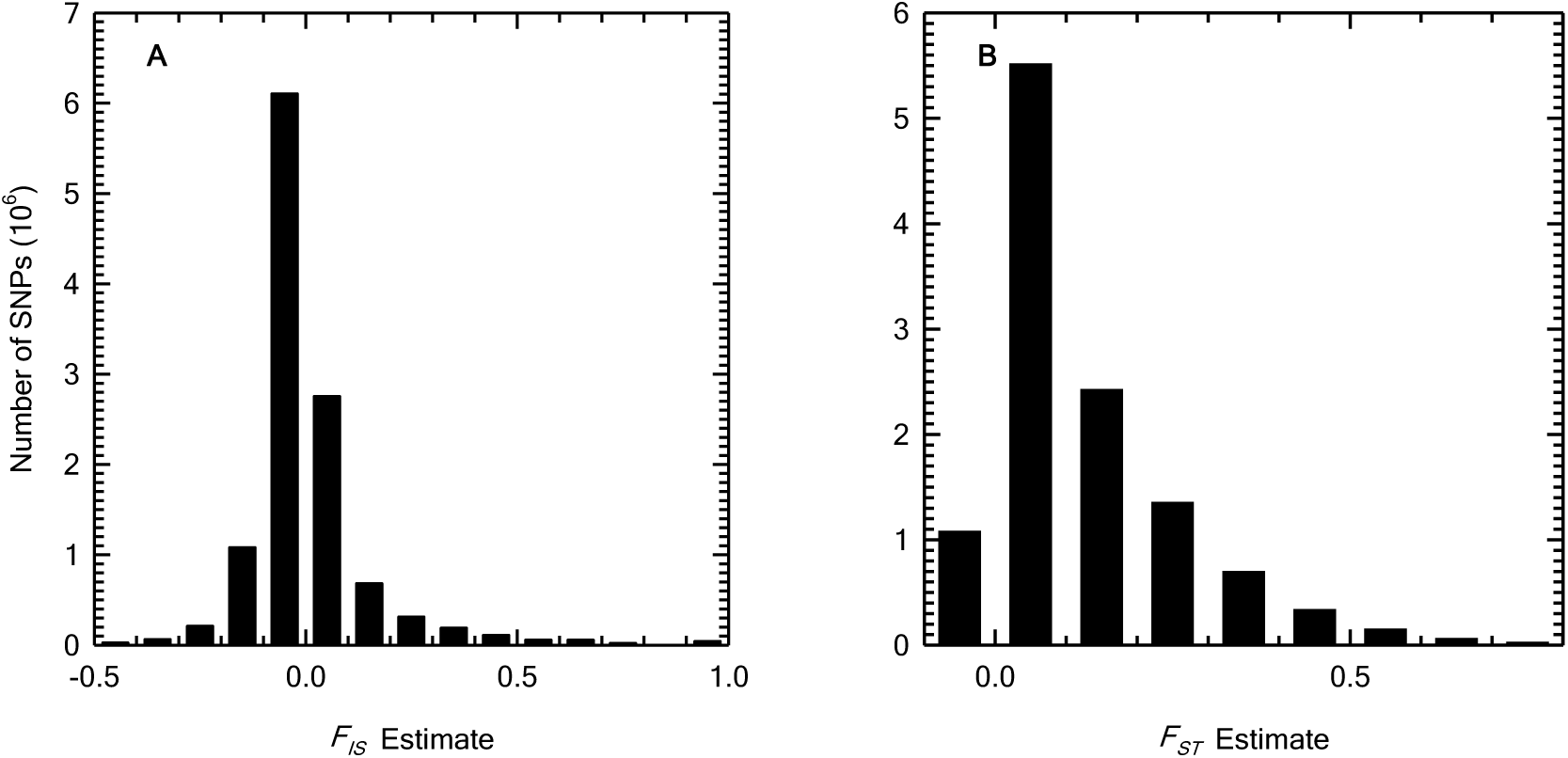
Genome-wide distribution of the *F_IS_* and *F_ST_* estimates. The distributions of the *F_IS_* (A) and *F_ST_* (B) estimates are shown using bins of size 0.1. The results are confined to sites with significant polymorphisms at the 5% level. The medians of the *F_IS_* and *F_ST_* estimates are -0.01 and 0.08, respectively.

**TABLE 3.** Wright’s (1951) fixation indices at genomic SNP sites. The mean is calculated using only sites with significant polymorphisms at the 5% level in at least one of the populations. SE denote the standard error of the mean.

| Estimate | Mean | SE | Number of SNPs |
| --- | --- | --- | --- |
| $\hat{F}_{IT}$ | 0.1280 | 0.00005 | 11,731,566 |
| $\hat{F}_{ST}$ | 0.1259 | 0.00004 | 11,731,566 |
| $\hat{F}_{IS}$ | 0.0019 | 0.00004 | 11,730,849 |

**TABLE 4.** F_ST_ estimates at sites in different functional categories. The mean is calculated using sites with significant polymorphisms at the 5% level in at least one of the populations. SE denotes the standard error of the mean. Intergenic sites are outside of untranslated regions (UTR), exons, and introns. Silent, replacement, and intron sites are, respectively four-fold redundant, zero-fold redundant and restricted intron (internal positions 8 to 34 from both ends; Lynch *et al*. 2017) sites.

| Category | Mean | SE | Number of SNPs |
| --- | --- | --- | --- |
| Silent | 0.1458 | 0.00020 | 526,078 |
| Replacement | 0.1082 | 0.00017 | 660,423 |
| Intron | 0.1473 | 0.00018 | 692,037 |
| 5'-UTR | 0.1305 | 0.00031 | 208,276 |
| 3'-UTR | 0.1270 | 0.00025 | 310,799 |
| Intergenic | 0.1213 | 0.00005 | 7,128,373 |

To examine the relationship between the sampling location and genetic differentiation, we built a neighbor-joining tree (Saitou and Nei 1987) of the mean pairwise *F_ST_* estimates (Figure 4A). Geographically close populations cluster together in the tree, indicating that gene flow among the populations is limited by the geographic distance. To further examine this hypothesis, we made a scatter plot of the geographic distance and mean pairwise *F_ST_* estimates at replacement and silent sites in protein-coding sequences (Figure 4B). The latter is positively correlated with the former (*p <* 0.05), supporting the hypothesis of distance-limited gene flow. *F_ST_* estimates are lower at replacement than silent sites, again likely because purifying selection decreases *F_ST_* estimates.

**Figure 4.**
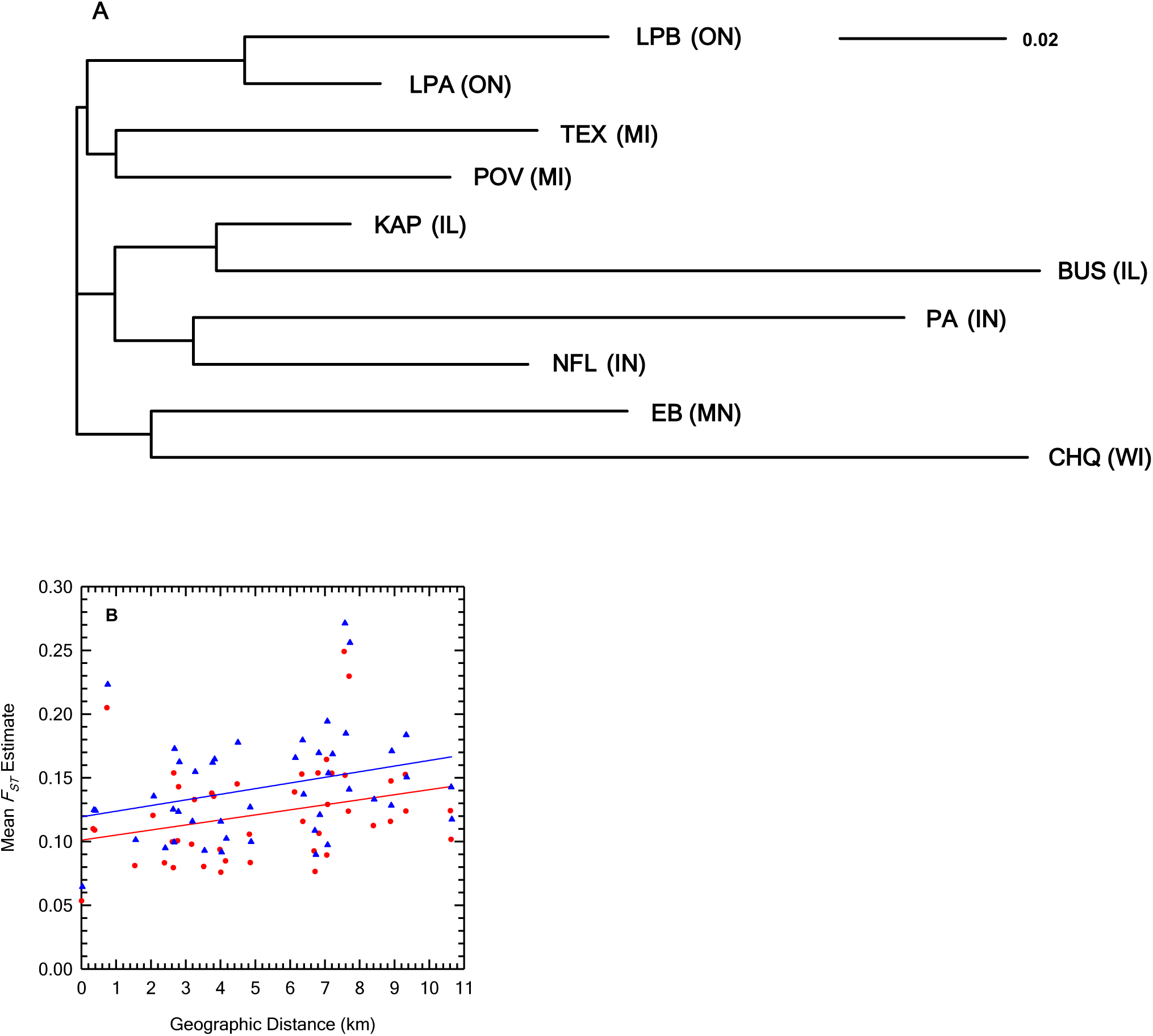
Pairwise *F_ST_* estimates. The neighbor-joining tree (A) and relationship with the geographic distance at replacement (red) and silent (blue) sites (B) are shown for the pairwise *F_ST_* estimates. The letters in parentheses in A denote the population state abbreviation. The scale bar in A is for the *F_ST_* estimates.

Because fixation-index estimates measured at individual SNP sites are highly variable (Weir and Hill 2002) and dependent on heterozygosity estimates (Nei 1973; Hedrick 1999; Maruki *et al*. 2012; Jakobbson *et al*. 2013; Alcala and Rosenberg 2017), we carried out sliding-window analyses to examine their spatial patterns along the scaffolds and to identify signatures of natural selection. Because *F_IS_* and *F_ST_* estimates are useful for finding signatures of natural selection (Black *et al*. 2001), we statistically identified their top and bottom outlier windows using the bootstrap at the 5% significance level. This revealed interesting peaks and valleys of the *F_IS_* estimates (Figure 5A), which may imply positive and balancing selection, respectively.

**Figure 5.**
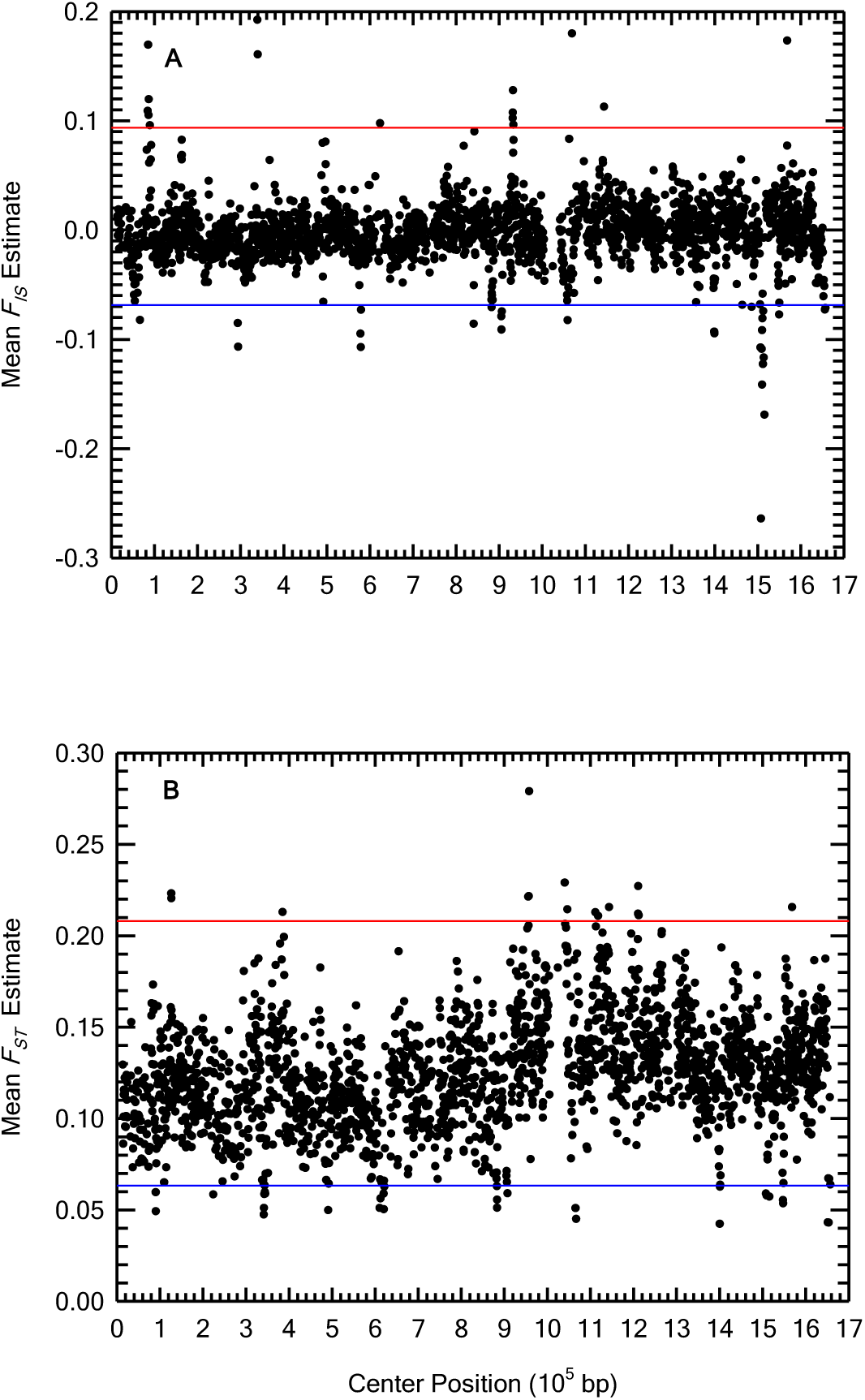
Spatial patterns of the *F_IS_* (A) and *F_ST_* (B) estimates on the largest scaffold. The red and blue lines denote the top and bottom critical values at the 5% significance level with the Bonferroni correction for multiple testing.

To infer biological mechanisms underlying the valleys of *F_IS_* estimates, we examined the contribution of the non-male producing (NMP) marker positions (Reisser *et al*. 2017) to the bottom *F_IS_* outlier regions at the 5% significance level. We recently identified 132 candidate NMP marker positions in an about 1.1 Mb region on chromosome I (Ye *et al*. in prep.), where all NMP clones are heterozygotes for a large number of markers, leading to an excess of heterozygotes compared to the Hardy-Weinberg equilibrium expectation. The *F_IS_* estimates at the NMP marker positions are highly negative (the mean = -0.189 with the standard error = 0.0025). However, only seven of 3,593 bottom *F_IS_* outlier regions contain the NMP marker positions (Table S2), showing that NMP marker positions are not the main cause of the bottom *F_IS_* outliers. In fact, the frequency of the NMP clones is generally low in the study populations. Unidentified biological mechanisms or balancing selection might be responsible for the other bottom *F_IS_* outliers, although care should be taken when invoking biological mechanisms underlying negative *F_IS_* values as they may result from mismapping or some other cryptic data issues (Maruki and Lynch 2015).

We also found interesting peaks and valleys for the *F_ST_* estimates (Figure 5B), which may imply local adaptation and purifying selection, respectively. By combining overlapping outlier windows, we made lists of top and bottom *F_IS_* and *F_ST_* outlier regions at the 5% significance level (Tables S1 to S4). Only seven of 2,491 top *F_ST_* outlier regions show significantly low mean heterozygosity within populations *π_S_* estimates (Table S3), indicating that the vast majority of the top *F_ST_* outlier regions result from high among-population genetic differentiation instead of low *π_S_*. To further study the relationship between heterozygosity and *F_ST_* estimates, we examined the relationship among *π_S_*, heterozygosity in the total population *π_T_*, and *F_ST_* estimates at replacement and silent sites in protein-coding sequences (Figures S1A, S1B, S2A, and S2B). *F_ST_* estimates generally increase with increased *π_S_* or *π_T_* estimates. This is mainly because the maximum possible value of *F_ST_* estimates increases with increased minor-allele frequency estimates in the total population at biallelic SNP sites (Maruki *et al*. 2012; Jakobbson *et al*. 2013; Alcala and Rosenberg 2017). Furthermore, *π_S_*, *π_T_*, and among-population heterozygosity *φ* estimates are significantly higher in top *F_ST_* outlier genes than in non-outlier genes at both replacement and silent sites (*p <* 0.05) (TABLE 5), which shows that the top *F_ST_* outlier genes result from increased among-population heterozygosity instead of decreased within-population heterozygosity.

**TABLE 5.**
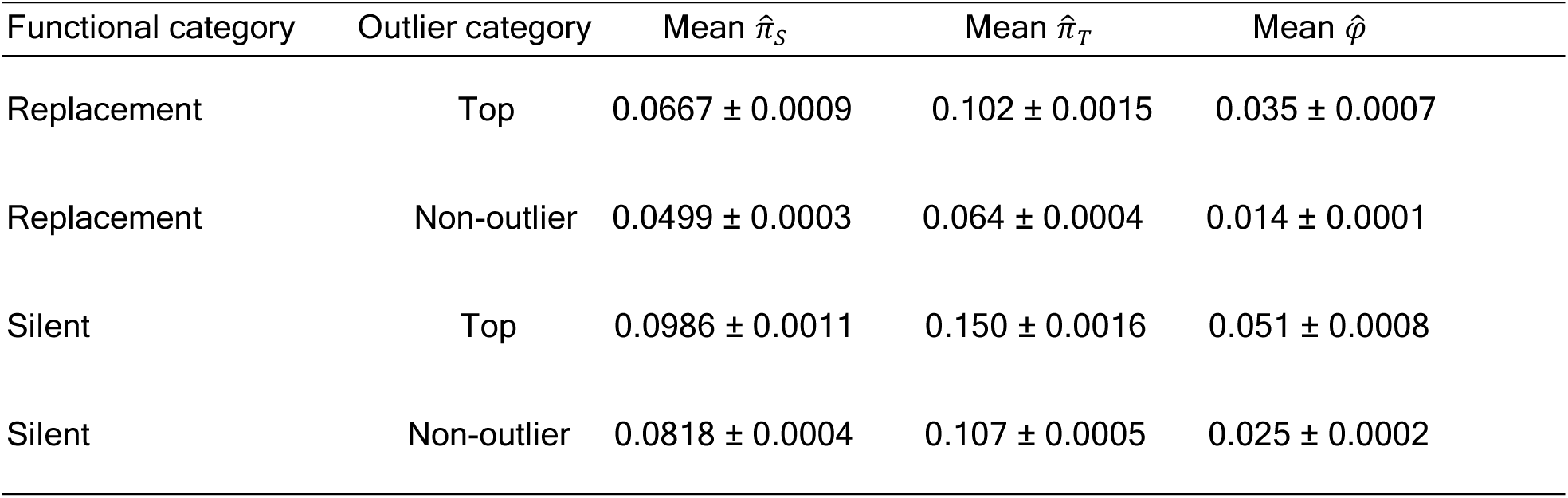
Comparison of the heterozygosity estimates for top *F_ST_* outlier genes and non-outlier genes in the genome. The mean and standard error of the mean heterozygosity within populations *π*_*S*_, heterozygosity in the total population *π*_*T*_, and among-population heterozygosity *φ* estimates at replacement and silent sites in top *F_ST_* outlier genes at the 5% significance level and non-outlier genes are shown. The mean is calculated using only sites with significant polymorphisms at the 5% level in at least one of the populations.

To infer the potential biological causes underlying the top *F_ST_* outlier genes, we carried out an enrichment analysis of the gene ontology (GO) terms in the top *F_ST_* outlier genes. Interestingly, there is striking enrichment of GO terms involved in food digestion (Hasler 1935; Schwerin *et al*. 2009; Schwarzenberger *et al*. 2012; Koussoroplis *et al*. 2017; Schwarzenberger and Flink 2018) in the top *F_ST_* outlier genes (TABLE 6 and Table S5), suggesting that environmental and/or nutritional differences among different populations may play an important role in the local adaptation of *Daphnia pulex*. In particular, we found strong enrichment of serine-type peptidases including trypsins and chymotrypsins (Table S6), which are major digestive enzymes in *Daphnia* (Von Elert *et al*. 2004; Schwerin *et al*. 2009; Schwarzenberger *et al*. 2012; Dӧolling *et al*. 2016). In stark contrast to this pattern, a completely different set of GO terms involved in cell structure are enriched in the bottom *F_ST_* outlier genes (TABLE 6 and Table S5), indicating genes maintaining cell structure are under especially strong purifying selection.

**TABLE 6.** Gene ontology terms enriched in the *F_ST_* outlier genes. Gene ontology (GO) terms belonging to the biological process (BP) or molecular function (MF) category with false-discovery rates (FDRs) < 0.05 are shown. The enrichment here is on a particular GO term in the *F_ST_* outlier genes at the 5% significance level compared to non-outlier genes.

| GO Term | Category | Outlier Type | FDR |
| --- | --- | --- | --- |
| peptidase activity | MF | Top | < 0.001 |
| hydrolase activity | MF | Top | < 0.01 |
| transmembrane transporter activity | MF | Top | < 0.05 |
| catalytic activity | MF | Top | < 0.05 |
| cell junction organization | BP | Bottom | < 0.001 |
| protein folding | BP | Bottom | < 0.01 |

To consider the confounding effect of the heterogeneity in the background genetic differentiation among different population pairs on identification of *F_ST_* outliers, we estimated *X^T^ X* (Günther and Coop 2013), which is an *F_ST_* analog taking heterogeneity in the background differentiation level into account, and carried out similar analyses. Although there are some differences, *X^T^ X* identified top and bottom outlier regions (Tables S7 and S8) similar to those identified using *F_ST_*. Supporting our results based on *F_ST_*, there is striking enrichment of GO terms involved in food digestion in the top *X^T^ X* outlier genes (Table S9), whereas those enriched in the bottom *X^T^X* outlier genes are involved in cell structure (Table S9). The same conclusions with *X^T^X* as those with *F_ST_* suggest that the heterogeneity in genetic differentiation among study populations is not large enough to significantly affect the identification of outlier regions.

To infer functional genetic changes underlying natural selection shaping the genetic differentiation among populations, we compared the mean *F_ST_* estimates over replacement and silent sites in each gene. Interestingly, genes showing a significant excess of amino-acid altering differentiation relative to silent-site differentiation (Table S10) include top *F_ST_* outlier genes coding proteins involved in food digestion. Also, many top *F_ST_* outlier genes showing a significant excess of amino-acid altering differentiation relative to silent-site differentiation do not have orthologs in the NCBI non-redundant protein database, indicating genes specific to *D. pulex* or a larger phylogenetic group of organisms encompassing *D. pulex* (*e.g.*, crustaceans) may play an important functional role in local adaptation. In contrast, the vast majority of genes showing a significant deficit of amino-acid altering differentiation relative to silent-site differentiation (Table S11), indicating purifying selection, have orthologs in the database.

To infer dominant selective forces shaping functional genomic changes of *D. pulex*, we calculated mean ratios of genetic changes over populations per replacement and silent sites over genes at three levels (*d′_N_/d′_S_* between species, *φ_N_/φ_S_* among populations, and *π_N_/π_S_* within populations). All of the mean ratios are significantly less than one (*p <* 0.05), and they decrease with higher level of the genetic differences (*π_N_/π_S_ > φ_N_/φ_S_ > d′_N_/d′_S_*) (Figure 6), indicating that purifying selection is the predominant average force in the long-term evolution of protein-coding sequences. Consistent with this, the mean of the analog of the neutrality index (NI) (Rand and Kann 1996), which is *π_N_/π_S_* divided by *d′_N_/d′_S_*, over genes is significantly greater than one (*p <* 0.05), and the mean of the analog of the direction of selection (DoS) (Stoletzki and Eyre-Walker 2011), which is *d′_N_/*(*d′_N_*+ *d′_S_*) *− π_N_/*(*π_N_* + *π_S_*), over genes is significantly less than zero (*p <* 0.05).

**Figure 6.**
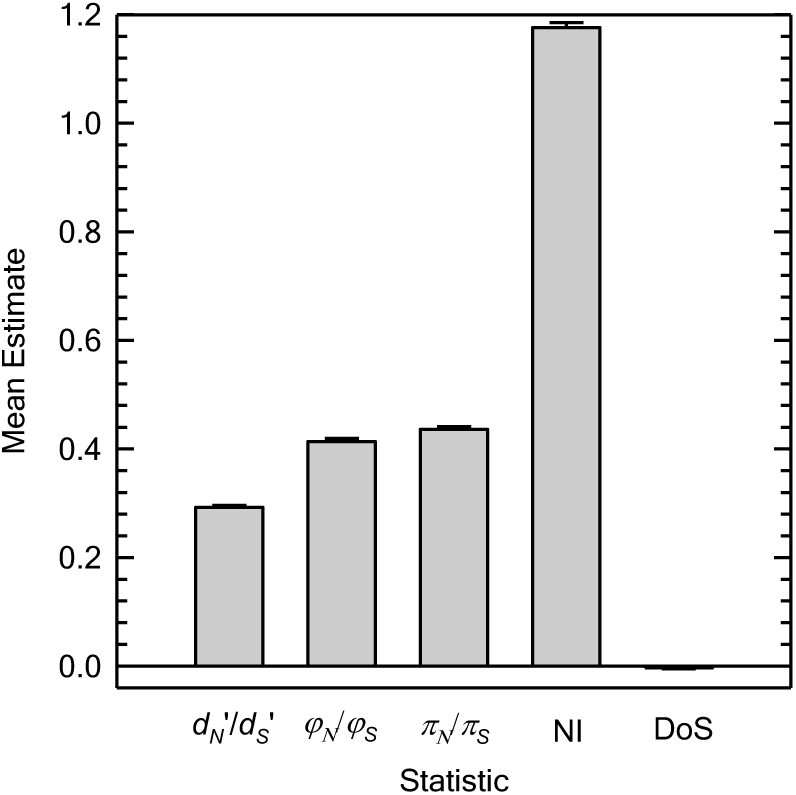
Comparison of genetic changes at replacement and silent sites in protein-coding sequences. Means ratios of genetic changes per replacement and silent sites over genes at three levels (*d*′*_N_*/*d*_*S*_′ between species, *φ*_*N*_/*φ*_*S*_ among populations, and *π*_*N*_/*π*_*S*_ within populations) are shown. Neutrality index (Rand and Kann 1996) and direction of selection (Stoletzki and Eyre-Walker 2011) estimates calculated from *d*_*N*_′, *d*_*S*_′, *π*_*N*_, *π*_*S*_ estimates are also shown.

To identify specific candidate genes under natural selection, we focused on genes that show signatures of positive or purifying selection in at least two of *d_N_ − d_S_*, neutrality index, and direction-of-selection estimates, finding 3,789 genes potentially under positive selection (Table S12). Interestingly, gene ontology terms enriched in the candidate genes (TABLE 7 and Table S13) are involved in food digestion, extending the importance of food for adaptations in *Daphnia* over a longer time scale. We also found 3,586 genes potentially under purifying selection (Table S14), where we found enrichment of gene ontology terms involved in cell structure and food digestion (TABLE 8 and Table S15).

**TABLE 7.** Gene ontology terms enriched in candidate genes under positive selection. Gene ontology (GO) terms belonging to molecular function terms with false-discovery rates (FDRs) < 0.05 are shown. The enrichment here is on a particular GO term in the candidate genes under positive selection with FDRs < 0.05 compared to non-outliers.

| GO Term | FDR |
| --- | --- |
| transmembrane transporter activity | < 0.001 |
| substrate-specific transmembrane transporter activity | < 0.001 |
| transporter activity | < 0.001 |
| catalytic activity | < 0.01 |
| substrate-specific transporter activity | < 0.01 |
| oxidoreductase activity | < 0.05 |

**TABLE 8.** Gene ontology terms enriched in candidate genes under purifying selection. Gene ontology (GO) terms belonging to molecular function terms with false-discovery rates (FDRs) < 0.05 are shown. The enrichment here is on a particular GO term in the candidate genes under purifying selection with FDRs < 0.05 compared to non-outliers.

| GO Term | FDR |
| --- | --- |
| binding | < 0.001 |
| ion binding | < 0.001 |
| RNA binding | < 0.001 |
| catalytic activity | < 0.001 |
| ligase activity | < 0.001 |
| nucleic acid binding | < 0.001 |
| enzyme binding | < 0.001 |
| protein binding | < 0.001 |
| transferase activity | < 0.001 |
| translation factor activity, RNA binding | < 0.01 |
| kinase activity | < 0.01 |
| organic cyclic compound binding | < 0.01 |
| heterocyclic compound binding | < 0.01 |
| transferase activity, transferring phosphorus-containing groups | < 0.01 |
| phosphatase activity | < 0.05 |
| phosphoric ester hydrolase activity | < 0.05 |
| cytoskeletal protein binding | < 0.05 |
| transferase activity, transferring acyl groups | < 0.05 |

To further study the signatures of natural selection in protein-coding sequences, we examined the relationship among *π_N_/π_S_*, *d_N_/d_S_*, and *φ_N_/φ_S_* estimates (Figures 7A, 7B, and 7C). Under neutral evolution, *d_N_/d_S_* and *φ_N_/φ_S_* are expected to be equal to *π_N_/π_S_*. The regression of *d_N_/d_S_* on *π_N_/π_S_* estimates shows that *d_N_/d_S_* estimates are greater than *π_N_/π_S_* estimates when *π_N_/π_S_* estimates are small, and *π_N_/π_S_* estimates are greater than *d_N_/d_S_* estimates when *π_N_/π_S_* estimates are large (Figure 7A). *d_N_/d_S_ > π_N_/π_S_* with small *π_N_/π_S_* indicates that positive directional selection tends to be strong and plays an important role in decreasing amino-acid altering polymorphisms relative to silent polymorphisms in the genome (Bazykin and Kondrashov 2011). *π_N_/π_S_ > d_N_/d_S_* with large *π_N_/π_S_* indicates that purifying selection tends to be weak (Ohta 1973) and does not efficiently eliminate amino-acid altering polymorphisms. Interestingly, the regression of *φ_N_/φ_S_* on *π_N_/π_S_* estimates shows an opposite pattern (Figure 7B). *φ_N_/φ_S_ < π_N_/π_S_* with small *π_N_/π_S_* indicates that purifying selection acting among populations tends to be strong and efficiently eliminates amino-acid altering polymorphisms to prevent fixation of deleterious alleles. *φ_N_/φ_S_ > π_N_/π_S_* with large *π_N_/π_S_* indicates that local adaptations tend to be ongoing and increase amino-acid altering polymorphisms. The slope of the regression of *d_N_/d_S_* on *φ_N_/φ_S_* estimates (Figure 7C) is lower than that in the regression of *d_N_/d_S_* on *π_N_/π_S_* estimates (Figure 7A), indicating that local adaptations significantly increase among-population amino-acid altering polymorphisms.

**Figure 7.**
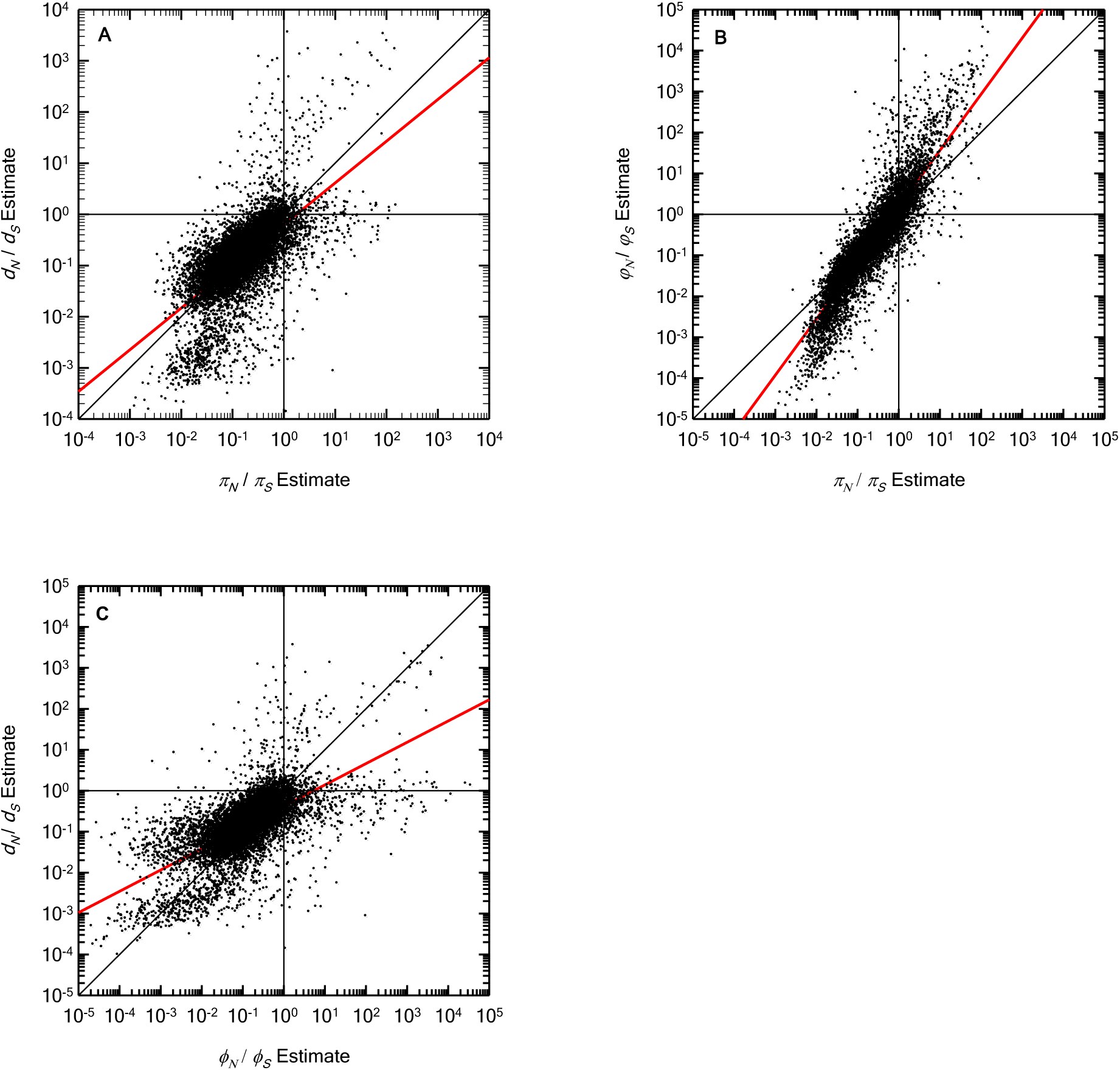
The relationship among *π*_*N*_/*π*_*S*_, *d*_*N*_/*d*_*S*_, and *φ*_*N*_/*φ*_*S*_ estimates in protein-coding sequences. Each dot denotes a gene. *π*_*N*_/*π*_*S*_, *d*_*N*_/*d*_*S*_,and *φ*_*N*_/*φ*_*S*_ are expected to be equal to one under neutrality (vertical and horizontal black lines). They are also expected to be equal to each other under neutrality (black diagonal lines). The 1) top left, 2) top right, 3) bottom right, and 4) bottom left quadrants in A indicate that the gene is under 1) positive directional selection, 2) positive directional selection (above the diagonal) or balancing selection (below the diagonal), 3) balancing selection, and 4) purifying selection (below the diagonal) or positive directional selection (above the diagonal), respectively. The 1) top left, 2) top right, 3) bottom right, and 4) bottom left quadrants in B indicate that the gene is under 1) local adaptation, 2) local adaptation (above the diagonal) or balancing selection (below the diagonal), 3) balancing selection, and 4) purifying selection (below the diagonal) or local adaptation (above the diagonal), respectively. The red lines are regression lines.

## DISCUSSION

This study represents the first genome-wide study of inter-population genetic variation in *Daphnia pulex* and one of the few to be performed in any species. The degree of genetic differentiation among *D. pulex* populations is moderate (mean 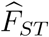 = 0.13) and defined to a large extent by the geographic distance between populations. It is somewhat higher than that in corresponding studies in humans (mean pairwise 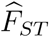 between populations in different continents ranging from 0.052 to 0.083) (The 1000 Genomes Project Consortium 2010) and flies (mean pairwise 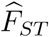 between populations within Africa ranging from 0.008 to 0.214) (Pool *et al*. 2012). We found interesting signatures of natural selection shaping the functional changes in the genome of *D. pulex*. In particular, the enrichment of genes involved in food digestion in the top *F_ST_* outlier genes indicates that environmental/nutritional differences among populations may play an important role in local adaptation of *D. pulex*.

Compared to the *F_ST_* estimates in most of the previous studies on genetic differentiation among populations in *D. pulex* (Crease *et al*. 1990; Lynch and Crease 1990; Lynch and Spitze 1994; Morgan *et al*. 2001; Allen *et al*. 2010), our *F_ST_* estimate is much lower. Because our estimate is based on genomic SNP data consisting of a large number of individuals generated by high-throughput sequencing, it is much less noisy compared to those in previous studies based on small numbers of molecular markers and individuals. Similar to our finding, a recent study on the population structure in *D. magna* using restriction-site associated DNA (RAD) sequencing (Orsini *et al*. 2013) reported a moderate amount of genetic differentiation among populations (mean *F_ST_* estimate of 0.12) based on SNP data. Our moderate *F_ST_* estimate among populations is in line with the potentially high dispersal capability of *Daphnia* (Havel and Shurin 2004; Figuerola *et al*. 2005). Assuming a simple island model (Wright 1931) and using the mean *F_ST_* estimate over silent and restricted intron SNP sites, a crude estimate of the product of the effective population size (*N_e_*) and migration rate (*m*) *N_e_m* is about 1.5. The relatively low genetic differentiation among populations in *D. pulex* is encouraging for population-genomic analyses in this species because top *F_ST_* outliers can be confidently identified only when the background differentiation level is low (Hoban *et al*. 2016). The positive correlation between the *F_ST_* estimates and geographic distance between populations in this study indicates that geographic distance plays an important role in limiting gene flow among populations, which is consistent with findings in some earlier studies (Crease *et al*. 1990; Lynch and Spitze 1994). This finding is also consistent with empirical evidence that historical dispersal of *Daphnia* has been rapid over short geographic distances and limited over large geographic distances (Havel and Shurin 2004). Furthermore, a recent study using RAD sequencing found the importance of geography in shaping the genetic structure of populations in *D. magna* (Fields *et al*. 2015).

The striking enrichment of genes involved in food digestion among the top *F_ST_* outlier genes provides a novel insight into the evolution of *D. pulex*. *Daphnia* often compete for a limited food supply, and reproduction is often limited by availability of food (Hebert 1978; Lynch 1989; Gliwicz 1990; Vanni and Lampert 1992; Tessier *et al*. 2000). The higher genetic differentiation among populations at replacement than that at silent sites in some of the top *F_ST_* outlier genes involved in food digestion (Table S9) suggests underlying adaptive changes, providing fuel for future hypothesis testing. Using RAD sequencing and mapping the sequence reads to the TCO assembly (Colbourne *et al*. 2011), Muñoz *et al*. (2016) also found that many of the genes showing significant genetic differentiation among five lake populations differing in trophic status are involved in metabolic processes in the sister species *D. pulicaria*. They also found many top *F_ST_* outlier genes involved in regulatory processes. Their similar finding in the sister species supports the importance of the nutritional differences among populations for local adaptation in *D. pulex*. In contrast to Muñoz *et al*. (2016), who intentionally selected diverse ecological settings, our *D. pulex* populations were sampled regardless of particular environmental variables, and yet still revealed overwhelming enrichment of genes involved in food digestion among the top *F_ST_* outlier genes. Therefore, our results not only provide an unbiased estimate of the genetic differentiation among populations, but affirm the importance of differences in food quality/quantity among populations as drivers of the evolution of *D. pulex* in general. Although other large-scale population-genomic studies in animals such as humans (The 1000 Genomes Project Consortium 2015) and flies (Langley *et al*. 2012; Pool *et al*. 2012) reported some genes involved in food digestion in top *F_ST_* outlier genes, the overwhelming enrichment of genes involved in food digestion among the top *F_ST_* outlier genes in *D. pulex* is notable.

Despite the existence of signatures of positive selection, the comparisons of the divergence and heterozygosity estimates between replacement and silent sites in protein-coding sequences revealed that purifying selection is the predominant force in the long-term evolution of protein-coding sequences in the *Daphnia* species. This result is consistent with our previous study of a single *D. pulex* population (Lynch *et al*. 2017), where we found predominant signatures of purifying selection preventing fixations of deleterious alleles at replacement sites. Taking advantage of data in multiple populations, we identified specific candidate genes under positive or purifying selection in this study. Gene ontology terms involved in food digestion are also enriched in genes showing signatures of positive selection in the long-term evolution, supporting the importance of food for adaptations in *Daphnia*.

This is one of the largest population-genomic studies ever performed in any organism, and establishes *D. pulex* as an excellent organism for carrying out population-genomic analyses to investigate mechanisms of evolution. As silent and restricted intron sites are essentially under neutral evolution (Lynch *et al*. 2017), they provide powerful means for inferring population demography/structure. In particular, we can use silent sites as the neutral standard for identifying signatures of natural selection in protein-coding sequences. The relatively large and historically stable effective sizes (Lynch *et al*. in prep.) of the study populations further help in identifying signatures of natural selection. Moreover, the homogeneous recombinational landscape (Lynch *et al*. in prep.) minimizes the confounding effect of heterogeneous recombination rates on signatures of selective sweeps (O’Reilly *et al*. 2008). Unlike isolates used in other large-scale population-genomic studies in *Drosophila* (Langley *et al*. 2012; Mackay *et al*. 2012) and *Arabidopsis* (Alonso-Blanco *et al*. 2016), our *Daphnia* isolates are not inbred. By analyzing the natural genotypes from the field, we showed that the study populations are essentially in Hardy-Weinberg equilibrium, which is often assumed in population-genetic analyses. As *Daphnia* can reproduce clonally, we can maintain the original genotypes in the laboratory for future experimental investigations for hypothesis testing, which is not possible in sexually reproducing organisms. These advantages and moderate genetic differentiation among study populations establish *D. pulex* as an excellent system for identifying targets of natural selection and investigating genotype-phenotype relationships in natural contexts.

## Acknowledgments

We thank Ken Spitze, Sen Xu, Jeffry Dudycha, and the Pfrender laboratory for sample collection. We also thank Emily Williams for maintaining *Daphnia* and James Ford for help in processing sequencing data. This work was supported by National Institutes of Health grants NIH-NIGMS R01-GM101672 and NIH-NIGMS R35-GM122566-01 and National Science Foundation grant DEB-1257806. The computations in this work were supported by the National Center for Genome Analysis Support, funded by National Science Foundation grant DBI-1458641 to Indiana University, and by Indiana University Research Technology’s computational resources. This work used the Extreme Science and Engineering Discovery Environment (XSEDE) (Towns *et al*. 2014), which is supported by National Science Foundation grant number ACI-1548562. Specifically, it used the Bridges system (Nystrom *et al*. 2015), which is supported by NSF award number ACI-1445606, at the Pittsburgh Supercomputing Center (PSC).

## APPENDIX

### Variance of the neutrality index and direction of selection

Assuming the divergence and heterozygosity estimates are uncorrelated and using the delta method (Lynch and Walsh 1998), the variance of the neutrality index (NI) is

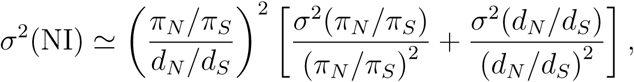

where *σ*^2^(*π_N_/π_S_*) and *σ*^2^(*d_N_/d_S_*) are found using the delta method and assuming that estimates at replacement and silent sites are uncorrelated as

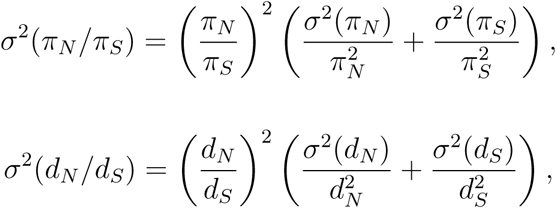

respectively. Similarly, the variance of the direction of selection (DoS) is

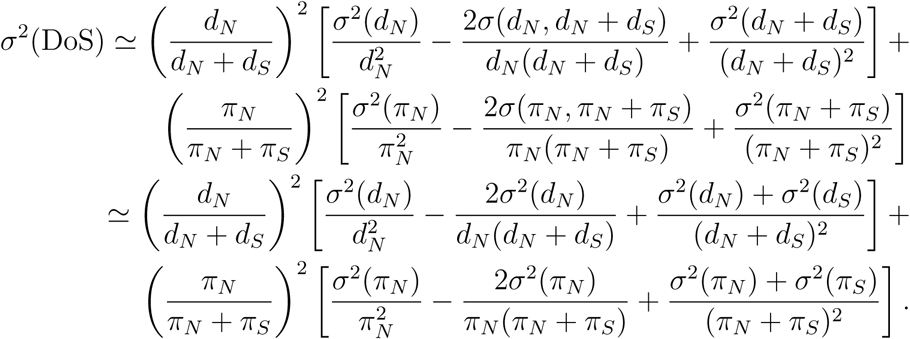

## LITERATURE CITED

Ackerman, M. S., P. Johri, K. Spitze, S. Xu, T. G. Doak et al., 2017 Estimating seven coefficients of pairwise relatedness using population-genomic data. Genetics 206: 105–118.

Al-Shahrour, F., R. Diaz-Uriarte and J. Dopazo, 2004 FatiGO: a web tool for finding significant associations of Gene Ontology terms with groups of genes. Bioinformatics 20: 578–580.

Alcala, N., and N. A. Rosenberg, 2017 Mathematical constraints on *FST*: biallelic markers in arbitrarily many populations. Genetics 206: 1581–1600.

Allen, M. R., R. A. Thum and C. E. Caceres, 2010 Does local adaptation to resources explain genetic differentiation among *Daphnia* populations? Mol Ecol 19: 3076–3087.

Alonso-Blanco, C., J. Andrade, C. Becker, F. Bemm, J. Bergelson et al., 2016 1,135 Genomes Reveal the Global Pattern of Polymorphism in Arabidopsis thaliana. Cell 166: 481–491.

Altschul, S. F., W. Gish, W. Miller, E. W. Myers and D. J. Lipman, 1990 Basic local alignment search tool. J Mol Biol 215: 403–410.

Ashburner, M., C. A. Ball, J. A. Blake, D. Botstein, H. Butler et al., 2000 Gene ontology: tool for the unification of biology. The Gene Ontology Consortium. Nat Genet 25: 25–29.

Barreiro, L. B., G. Laval, H. Quach, E. Patin and L. Quintana-Murci, 2008 Natural selection has driven population differentiation in modern humans. Nat Genet 40: 340–345.

Bazykin, G. A., and A. S. Kondrashov, 2011 Detecting past positive selection through ongoing negative selection. Genome Biol Evol 3: 1006–1013.

Black, W. C. IV., C. F. Baer, M. F. Antolin and N. M. DuTeau, 2001 Population genomics: genome-wide sampling of insect populations. Annu Rev Entomol 46: 441–469.

Bolger, A. M., M. Lohse and B. Usadel, 2014 Trimmomatic: a flexible trimmer for Illumina sequence data. Bioinformatics 30: 2114–2120.

Breese, M. R., and Y. Liu, 2013 NGSUtils: a software suite for analyzing and manipulating next-generation sequencing datasets. Bioinformatics 29: 494–496.

Brendonck, L., and L. De Meester, 2003 Egg banks in freshwater zooplankton: evolutionary and ecological archives in the sediments. Hydrobiologia 491: 65–84

Camacho, C., G. Coulouris, V. Avagyan, N. Ma, J. Papadopoulos et al., 2009 BLAST+: architecture and applications. BMC Bioinformatics 10: 421.

Colbourne, J. K., M. E. Pfrender, D. Gilbert, W. K. Thomas, A. Tucker et al., 2011 The ecoresponsive genome of *Daphnia pulex*. Science 331: 555–561.

Conesa, A., S. Gotz, J. M. Garcia-Gomez, J. Terol, M. Talon et al., 2005 Blast2GO: a universal tool for annotation, visualization and analysis in functional genomics research. Bioinformatics 21: 3674–3676.

Crease, T. J., M. Lynch and K. Spitze, 1990 Hierarchical analysis of population genetic variation in mitochondrial and nuclear genes of *Daphnia pulex*. Mol Biol Evol 7: 444–458.

DePristo, M. A., E. Banks, R. Poplin, K. V. Garimella, J. R. Maguire et al., 2011 A framework for variation discovery and genotyping using next-generation DNA sequencing data. Nat Genet 43: 491–498.

Dӧlling, R., D. Becker, S. Hawat, M. Koch, A. Schwarzenberger et al., 2016 Adjustments of serine proteases of *Daphnia pulex* in response to temperature changes. Comp Biochem Physiol B Biochem Mol Biol 194-195: 1-10.

Efron, B., and R. Tibshirani, 1993 An introduction to the bootstrap. Chapman & Hall, New York.

Fields, P. D., C. Reisser, M. Dukic, C. R. Haag and D. Ebert, 2015 Genes mirror geography in *Daphnia magna*. Mol Ecol 24: 4521–4536.

Figuerola, J., A. J. Green, T. C. Michot, 2005 Invertebrate eggs can fly: evidence of waterfowl-mediated gene flow in aquatic invertabrates. The American Naturalist 165: 274–280.

Genomes Project, C., G. R. Abecasis, D. Altshuler, A. Auton, L. D. Brooks et al., 2010 A map of human genome variation from population-scale sequencing. Nature 467: 1061–1073.

Genomes Project, C., A. Auton, L. D. Brooks, R. M. Durbin, E. P. Garrison et al., 2015 A global reference for human genetic variation. Nature 526: 68–74.

Gliwicz, Z. M., 1990 Food thresholds and body size in cladocerans. Nature 343: 638–640.

Günther, T., and G. Coop, 2013 Robust identification of local adaptation from allele frequencies. Genetics 195: 205–220.

Hasler, A. D., 1935 The physiology of digestion of plankton crustacea, I. some digestive enzymes of Daphnia. Biol. Bull. 68: 207–214

Havel, J. E., and J. B. Shurin, 2004 Mechanisms, effects, and scales of dispersal in freshwater zooplankton. Limnology and Oceanography 49: 1229–1238.

Hebert, P. D. N., 1978 The population biology of *Daphnia* (*Crustacea*, *Daphnidae*). Biological Reviews 53: 387–426

Hedrick, P. W., 1999 Perspective: highly variable loci and their interpretation in evolution and conservation. Evolution 53: 313–318.

Hijmans, R. J., 2017 geosphere: Spherical Trigonometry. R package version 1.5-7. https://CRAN.R-project.org/package=geosphere

Hoban, S., J. L. Kelley, K. E. Lotterhos, M. F. Antolin, G. Bradburd et al., 2016 Finding the genomic basis of local adaptation: pitfalls, practical solutions, and future directions. Am Nat 188: 379–397.

Hohenlohe, P. A., S. Bassham, P. D. Etter, N. Stiffler, E. A. Johnson et al., 2010 Population genomics of parallel adaptation in threespine stickleback using sequenced RAD tags. PLoS Genet 6: e1000862.

Innes, D., 1991 Geographic patterns of genetic differentiation among sexual populations of *Daphnia pulex*. Canadian Journal of Zoology 69: 995–1003

Jakobsson, M., M. D. Edge and N. A. Rosenberg, 2013 The relationship between *FST* and the frequency of the most frequent allele. Genetics 193: 515–528.

Jones, P., D. Binns, H. Y. Chang, M. Fraser, W. Li et al., 2014 InterProScan 5: genome-scale protein function classification. Bioinformatics 30: 1236–1240.

Jurka, J., V. V. Kapitonov, A. Pavlicek, P. Klonowski, O. Kohany et al., 2005 Repbase Update, a database of eukaryotic repetitive elements. Cytogenet Genome Res 110: 462–467.

Keith, N., A. E. Tucker, C. E. Jackson, W. Sung, J. I. Lucas Lledo et al., 2016 High mutational rates of large-scale duplication and deletion in *Daphnia pulex*. Genome Res 26: 60–69.

Kielbasa, S. M., R. Wan, K. Sato, P. Horton and M. C. Frith, 2011 Adaptive seeds tame genomic sequence comparison. Genome Res 21: 487–493.

Koussoroplis, A. M., A. Schwarzenberger and A. Wacker, 2017 Diet quality determines lipase gene expression and lipase/esterase activity in *Daphnia pulex*. Biol Open 6: 210–216.

Langley, C. H., K. Stevens, C. Cardeno, Y. C. Lee, D. R. Schrider et al., 2012 Genomic variation in natural populations of *Drosophila melanogaster*. Genetics 192: 533–598.

Li, H., B. Handsaker, A. Wysoker, T. Fennell, J. Ruan et al., 2009 The Sequence Alignment/Map format and SAMtools. Bioinformatics 25: 2078–2079.

Luikart, G., P. R. England, D. Tallmon, S. Jordan and P. Taberlet, 2003 The power and promise of population genomics: from genotyping to genome typing. Nat Rev Genet 4: 981–994.

Lynch, M., 1983 Ecological genetics of *Daphnia Pulex*. Evolution 37: 358–374.

Lynch, M., 1989 The life history consequences of resource depression in *Daphnia pulex*. Ecology 70: 246–256.

Lynch, M., and T. J. Crease, 1990 The analysis of population survey data on DNA sequence variation. Mol Biol Evol 7: 377–394.

Lynch, M., and K. Spitze, 1994 Evolutionary genetics of Daphnia. pp. 109–128. In L. Real (ed.) Ecological Genetics. Princeton Univ. Press.

Lynch, M., and Walsh, B. 1998 Genetics and analysis of quantitative traits. Sinauer Associates, Sunderland, Mass.

Lynch, M., R. Gutenkunst, M. Ackerman, K. Spitze, Z. Ye et al., 2017 Population genomics of *Daphnia pulex*. Genetics 206: 315–332.

Lynch, M., B. Haubold, P. Pfaffelhuber, and T. Maruki, Inference of historical population-size changes with allele-frequency data. (in prep.)

Lynch, M., Z. Ye, and T. Maruki, The recombinational landscape in Daphnia pulex. (in prep.)

Mackay, T. F. C., S. Richards, E. A. Stone, A. Barbadilla, J. F. Ayroles et al., 2012 The *Drosophila melanogaster* genetic reference panel. Nature 482: 173–178.

Maruki, T., S. Kumar and Y. Kim, 2012 Purifying selection modulates the estimates of population differentiation and confounds genome-wide comparisons across single-nucleotide polymorphisms. Mol Biol Evol 29: 3617–3623.

Maruki, T., and M. Lynch, 2015 Genotype-frequency estimation from high-throughput sequencing data. Genetics 201: 473–486.

Maruki, T., and M. Lynch, 2017 Genotype calling from population-genomic sequencing data. G3 (Bethesda) 7: 1393-1404.

McKenna, A., M. Hanna, E. Banks, A. Sivachenko, K. Cibulskis et al., 2010 The Genome Analysis Toolkit: a MapReduce framework for analyzing next-generation DNA sequencing data. Genome Res 20: 1297–1303.

Morgan, K. K., J. Hicks, K. Spitze, L. Latta, M. E. Pfrender et al., 2001 Patterns of genetic architecture for life-history traits and molecular markers in a subdivided species. Evolution 55: 1753–1761.

Muñoz, J., A. Chaturvedi, L. De Meester and L. J. Weider, 2016 Characterization of genome-wide SNPs for the water flea *Daphnia pulicaria* generated by genotyping-by-sequencing (GBS). Sci Rep 6: 28569.

Myhre, S., H. Tveit, T. Mollestad and A. Laegreid, 2006 Additional gene ontology structure for improved biological reasoning. Bioinformatics 22: 2020–2027.

Nei, M., 1973 Analysis of gene diversity in subdivided populations. Proc Natl Acad Sci U S A 70: 3321–3323.

Nystrom, N. A., M. J. Levine, R. Z. Roskies and J. R. Scott, 2015 Bridges: a uniquely flexible HPC resource for new communities and data analytics, pp. 1–8 in Proceedings of the 2015 XSEDE Conference: Scientific Advancements Enabled by Enhanced Cyberinfrastructure. ACM, St. Louis, Missouri.

Ohta, T., 1973 Slightly deleterious mutant substitutions in evolution. Nature 246: 96–98.

O’Reilly, P. F., E. Birney and D. J. Balding, 2008 Confounding between recombination and selection, and the Ped/Pop method for detecting selection. Genome Research 18: 1304–1313.

Orsini, L., J. Mergeay, J. Vanoverbeke and L. De Meester, 2013 The role of selection in driving landscape genomic structure of the waterflea *Daphnia magna*. Mol Ecol 22: 583–601.

Paradis, E., J. Claude and K. Strimmer, 2004 APE: Analyses of Phylogenetics and Evolution in R language. Bioinformatics 20: 289–290.

Pool, J. E., R. B. Corbett-Detig, R. P. Sugino, K. A. Stevens, C. M. Cardeno et al., 2012 Population genomics of sub-saharan *Drosophila melanogaster*: African diversity and non-African admixture. PLoS Genet 8: e1003080.

Rand, D. M., and L. M. Kann, 1996 Excess amino acid polymorphism in mitochondrial DNA: contrasts among genes from *Drosophila*, mice, and humans. Mol Biol Evol 13: 735–748.

Reisser, C. M. O., D. Fasel, E. Hurlimann, M. Dukic, C. Haag-Liautard et al., 2017 Transition from environmental to partial genetic sex determination in *Daphnia* through the evolution of a female-determining incipient W chromosome. Molecular Biology and Evolution 34: 575–588.

Saitou, N., and M. Nei, 1987 The neighbor-joining method: a new method for reconstructing phylogenetic trees. Mol Biol Evol 4: 406–425.

Schwarzenberger, A., C. J. Kuster and E. Von Elert, 2012 Molecular mechanisms of tolerance to cyanobacterial protease inhibitors revealed by clonal differences in *Daphnia magna*. Mol Ecol 21: 4898–4911.

Schwarzenberger, A., and P. Fink, 2018 Gene expression and activity of digestive enzymes of *Daphnia pulex* in response to food quality differences. Comp Biochem Physiol B Biochem Mol Biol 218: 23–29.

Schwerin, S., B. Zeis, T. Lamkemeyer, R. J. Paul, M. Koch et al., 2009 Acclimatory responses of the *Daphnia pulex* proteome to environmental changes. II. Chronic exposure to different temperatures (10 and 20 degrees C) mainly affects protein metabolism. BMC Physiol 9: 8.

Stoletzki, N., and A. Eyre-Walker, 2011 Estimation of the neutrality index. Mol Biol Evol 28: 63–70.

Storey, J. D., and R. Tibshirani, 2003 Statistical significance for genomewide studies. Proc Natl Acad Sci U S A 100: 9440–9445.

Tessier, A. J., M. A. Leibold, and J. Tsao, 2000 A fundamental trade-off in resource exploitation by *Daphnia* and consequences to planktonic communities. Ecology 81: 826–841

Towns, J., T. Cockerill, M. Dahan, I. Foster, K. Gaither et al.,, 2014. XSEDE: Accelerating Scientific Discovery. Computing in Science & Engineering. 16(5):62–74.

Tucker, A. E., M. S. Ackerman, B. D. Eads, S. Xu and M. Lynch, 2013 Population-genomic insights into the evolutionary origin and fate of obligately asexual *Daphnia pulex*. Proc Natl Acad Sci U S A 110: 15740–15745.

Van der Auwera, G. A., M. O. Carneiro, C. Hartl, R. Poplin, G. Del Angel et al., 2013 From FastQ data to high confidence variant calls: the Genome Analysis Toolkit best practices pipeline. Curr Protoc Bioinformatics 11: 11 10 11-11 10 33.

Vanni, M. J., and W. Lampert, 1992 Food quality effects on life history traits and fitness in the generalist herbivore *Daphnia*. Oecologia 92: 48–57.

von Elert, E., M. K. Agrawal, C. Gebauer, H. Jaensch, U. Bauer et al., 2004 Protease activity in gut of Daphnia magna: evidence for trypsin and chymotrypsin enzymes. Comp Biochem Physiol B Biochem Mol Biol 137: 287–296.

Weir, B. S., 1996 Genetic data analysis II: methods for discrete population genetic data. Sinauer Associates, Sunderland, Mass.

Weir, B. S., and C. C. Cockerham, 1984 Estimating F-Statistics for the analysis of population structure. Evolution 38: 1358–1370.

Weir, B. S., and W. G. Hill, 2002 Estimating F-statistics. Annu Rev Genet 36: 721–750.

Wright, S., 1931 Evolution in mendelian populations. Genetics 16: 97–159.

Wright, S., 1951 The genetical structure of populations. Ann Eugen 15: 323–354.

Xu, S., K. Spitze, M. S. Ackerman, Z. Ye, L. Bright et al., 2015 Hybridization and the origin of contagious asexuality in *Daphnia pulex*. Mol Biol Evol 32: 3215–3225.

Ye, Z., S. Xu, K. Spitze, J. Asselman, X. Jiang et al., 2017 A new reference genome assembly for the microcrustacean *Daphnia pulex*. G3 (Bethesda) 7: 1405-1416.

Ye, Z., C. Molinier, C. Zhao, C. R. Haag, and M. Lynch, Genetic control of male production in Daphnia pulex. (in prep.)

